# AAV-mediated *CSNK2B* gene replacement rescues ASD-relevant phenotypes and establishes EEG biomarkers for translation in *Csnk2b* haploinsufficient mice

**DOI:** 10.1101/2025.10.23.684260

**Authors:** Chaodong Ding, Xiaqing Wang, Yiting Yuan, Yuefang Zhang, Yixin Hu, Ailian Du, Zilong Qiu

## Abstract

*De novo CSNK2B* variants are strongly associated with autism spectrum disorder (ASD) and early-onset epilepsy, yet *in vivo* therapeutic evidence and translatable biomarkers remain limited. We generated *Csnk2b* haploinsufficient (*Csnk2b^+/–^*) mice that recapitulate core clinical features—ASD-like social and cognitive deficits, heightened anxiety, spontaneous seizures, cortical and hippocampal structural compromise, and reduced PV-interneuron density with impaired cortical inhibition. Neonatal brain-wide AAV-PHP.eB–mediated CSNK2B replacement (hsyn or CAG promoters, retro-orbital P3) restored cortical/hippocampal structure, normalized neuronal numbers, prolonged survival, and ameliorated seizures and ASD-like behaviors with comparable efficacy across promoters. Critically, treatment also corrected quantitative EEG signatures that are directly deployable in human trials: theta and gamma band power were normalized, theta/gamma (and beta/gamma) ratios recovered, and inter-areal coherence and gamma-band effective connectivity were restored, indicating re-established excitation/inhibition balance and large-scale network coordination. These EEG endpoints constitute a compact, noninvasive biomarker panel of target engagement and physiological efficacy that can be harmonized with clinical EEG to support dose finding, early decision-making, and longitudinal monitoring in *CSNK2B* gene-therapy trials. Together, our data provide preclinical proof-of-concept that early AAV-mediated CSNK2B replacement is therapeutically effective and nominate band-limited EEG metrics as translational readouts to accelerate first-in-human studies for CSNK2B-linked neurodevelopmental disorders.

## Introduction

In recent years, the link between *CSNK2B* (Casein kinase 2 beta) and multiple neurodevelopmental disorders has been increasingly recognized due to advancements in gene sequencing technologies. For instance, splice-site mutations (c.176-2A>G; c.292-2A>G) in this gene have been found in autistic children^1^. ASD, a complex neurodevelopmental disorder, is marked by impaired social interaction and repetitive behavior patterns^2^. Furthermore, heterozygous pathogenic mutations in this gene are also the main cause of Poirier-Bienvenu neurodevelopmental syndrome (POBINDS)^3–5^. Most POBINDS patients have *de novo CSNK2B* mutations, with clinical features like epileptic seizures, developmental delay, facial dysmorphism, and intellectual disability^6,7^. Epilepsy occurs in most *CSNK2B* mutation patients, with an early onset (usually before age two)^4,7–9^. It includes various types like myoclonic epilepsy, tonic-clonic epilepsy, and febrile seizures, and its severity correlates with the degree of neurodevelopmental defects^7,9^. So far, at least 25 cases of neurodevelopmental diseases associated with *CSNK2B* mutations have been reported in the literature^10^, yet how these mutations affect neurological function and behavioral manifestations remains unclear.

*CSNK2B* encodes the regulatory subunit of casein kinase II (CK2) and is more highly expressed in the human brain during the embryonic period than postnatally^11^, indicating its importance in brain development. CK2 is highly expressed in the brain and is involved in processes like signal transduction and transcription regulation^12,13^. For example, in rats with cerebral hemorrhage, CK2 can downregulate the expression of NMDA receptor (NR2B) through phosphorylation, thereby reducing neuronal apoptosis, inflammation, and oxidative stress^13^. Homozygous knockout of *Csnk2b* leads to weakened cell proliferation in mouse embryos and results in embryonic lethality^14^. Moreover, knocking out *Csnk2b* in embryonic neural progenitors can affect their proliferation and differentiation into oligodendrocytes^15^. In addition to ASD and POBINDS, *CSNK2B* has also been identified as a schizophrenia risk gene^11^. *CSNK2B* is low expressed in schizophrenia patients, and inhibiting *Csnk2b* expression can affect neuronal morphology and synaptic transmission function in mouse^11^. These findings highlight the important role of *Csnk2b* in neural development and function. However, there have been no reports on the investigation of ASD- and epilepsy-related phenotypes in *Csnk2b* knockout mice, along with the subsequent therapeutic interventions.

In this study, we employed CRISPR/Cas9 technology to generate a *Csnk2b* heterozygous knockout mouse model (*Csnk2b*^+/-^) aimed at investigating the impact of *Csnk2b* mutations on mouse behavior and neurodevelopment. Our results demonstrated that *Csnk2b*^+/-^ mice exhibited behavioral traits associated with ASD, such as social impairment, heightened anxiety, and spatial learning and memory deficits, along with epilepsy-related behaviors and electroencephalographic signals. Moreover, these mice experienced growth retardation, aberrant brain structures, and lower survival rates. Further analysis revealed that reduced *Csnk2b* expression disrupted the development of specific cortical neurons and neural network synchronization. Subsequently, we administered AAV-mediated gene replacement therapy to neonatal *Csnk2b*^+/-^ mice. This intervention remarkably enhanced their neurodevelopment, network synchronization, and survival rate. Notably, their ASD-like behaviors and epileptic symptoms were ameliorated. In summary, our findings provide significant experimental evidence, offering promising insights into potential therapeutic strategies for neurodevelopmental disorders linked to *CSNK2B* mutations.

## Results

### *Csnk2b* haploinsufficiency causes brain structural abnormalities, reduced survival rate, ASD-like behaviors, and epileptic symptoms in mice

Several studies have reported various *de novo* mutations in the *CSNK2B* gene in patients with intellectual disability, epilepsy, and ASD, including splicing mutations, nonsense mutations, and missense mutations (Figure 1A)^1,10,16,17^. To explore the functional effects of *CSNK2B* mutations, we first used the CRISPR/Cas9 system to knock out the *Csnk2b* gene in mice. After designing single-guide RNAs (sgRNAs) flanking exon 3 of the *Csnk2b*, we successfully deleted 841 base pairs in this region (Figures S1A and S1B). Subsequently, we detected the expression levels and found that the expression of *Csnk2b* in the brains of *Csnk2b*^+/-^ mice was reduced by approximately 50% compared to that in wild-type (WT) mice (Figures S1C-S1E).

**Figure 1.**
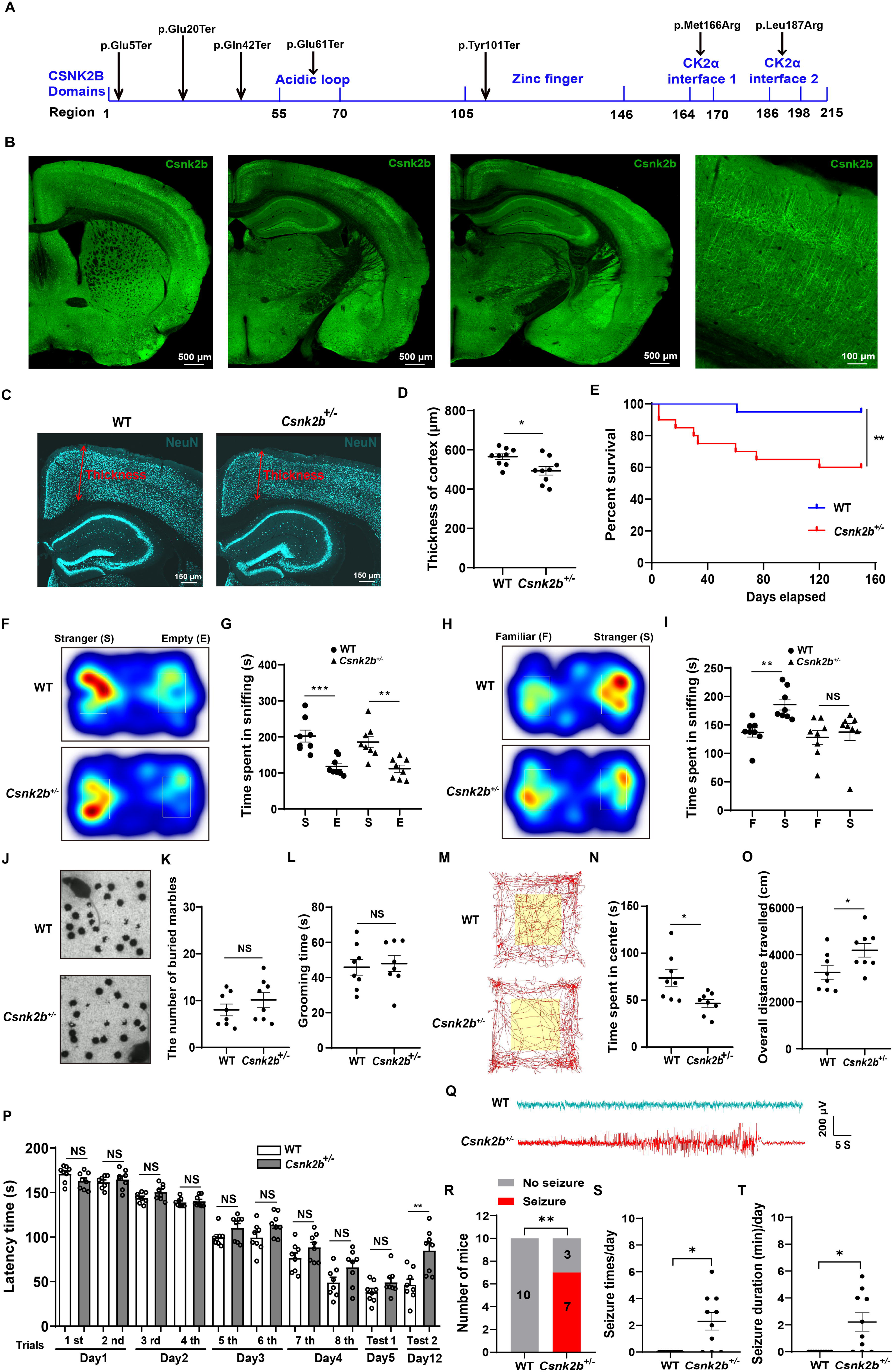
*Csnk2b* haploinsufficiency causes brain structural abnormalities, reduced survival rate, ASD-like behaviors, and epileptic symptoms in mice. (A) Diagrammatic representation of *de novo* mutations within the *CSNK2B* gene. (B) Immunofluorescence staining of *Csnk2b* in different brain regions of WT mouse brain sections. (C) Immunofluorescence staining of neurons in the cortex and hippocampus of WT and *Csnk2b*^+/-^ mice. (D) Quantification of cortical thickness from the staining shown in (C) (n = 9 slices from 3 mice). (E) Survival curves of WT and *Csnk2b*^+/-^ mice up to 150 days postnatal (n = 20 per group). (F) Heatmap tracing of WT and *Csnk2b*^+/-^ mice during the sociability test. (G) Time spent interacting with stranger mice versus empty cages (n = 8 per group). (H) Heatmap tracing of WT and *Csnk2b*^+/-^ mice during the social novelty test. (I) Quantification of social interaction time (n = 8 per group). (J) Representative images of the marble burying test results. (K) Number of marbles buried (n = 8 per group). (L) Self-grooming time in WT and *Csnk2b*^+/-^ mice (n = 8 per group). (M) Representative trajectories of WT and *Csnk2b*^+/-^ mice in the open field test. (N) Time spent in the central area (n = 8 per group). (O) Total distance traveled by WT and *Csnk2b*^+/-^ mice (n = 8 per group). (P) Time taken by WT and *Csnk2b*^+/-^ mice to locate the escape box in the Barnes maze test (n = 8 per group). (Q) Representative ECoG recordings showing epileptiform discharges in *Csnk2b*^+/-^ mice and absence of abnormal discharges in WT mice. (R) Quantification of the incidence of spontaneous epileptic seizures in *Csnk2b*^+/-^ mice and WT mice (n = 10 mice for each group). (S) Daily frequency of spontaneous seizures in *Csnk2b*^+/-^ mice and WT mice (n = 10 mice for each group). (T) Total daily duration of spontaneous seizures in *Csnk2b*^+/-^ mice and WT mice (n = 10 mice for each group).Data are expressed as mean±SD. Statistical analysis was performed using two-tailed Student’s t-test (D, G, I, K, L, N, O, P), log rank (Mantel-Cox) test (E), Fisher’s exact test (R), and Mann-Whitney U test (S, T). \**P <* 0.05, \*\**P <* 0.01, \*\*\**P <* 0.001, NS (not significant).

To investigate the role of *Csnk2b* in development, we measured the body weights of *Csnk2b*^+/-^ mice and WT mice at postnatal day 21 and day 60. The results showed that *Csnk2b*^+/-^ mice had significantly lower body weights at both time points compared to WT mice (Figure S1F). Additionally, the brain weights of *Csnk2b*^+/-^ mice were significantly lower than those of WT mice at these two stages (Figure S1G). Further examination revealed that the height and width of the cerebral cortex were significantly reduced in *Csnk2b*^+/-^ mice compared to WT mice (Figure S1H). Staining of mouse brain sections revealed that *Csnk2b* is expressed in multiple brain regions, including the cortex and hippocampus (Figure 1B). Within the cortical region, *Csnk2b* is highly expressed in the dendrites and axons of neurons (Figure 1B). Neuronal staining showed that the cortical thickness was decreased, the hippocampal structure was abnormal, and the lateral ventricles were enlarged in *Csnk2b*^+/-^ mice (Figures 1C and 1D). Finally, we found that haploinsufficiency of *Csnk2b* significantly reduced the survival rate of mice, with *Csnk2b*^+/-^ mice having a survival rate of only 60% at 120 days postnatal (Figure 1E).

To evaluate the social ability of *Csnk2b*^+/-^ mice, we first conducted the three-chamber test. During the sociability test, we found that *Csnk2b*^+/-^ mice exhibited similar social abilities to WT mice, spending more time interacting with a stranger mouse rather than exploring an empty cage (Figures 1F and 1G). However, when confronted both with a stranger and a familiar mouse, *Csnk2b*^+/-^ mice did not display the same social preference as WT mice, instead allocating approximately equal time to exploring both mice (Figures 1H and 1I).

Subsequently, we investigated the ability of *Csnk2b*^+/-^ mice to explore and recognize novel and familiar objects. When the right chamber was empty and the left chamber contained a novel object, *Csnk2b*^+/-^ mice, similar to WT mice, spent more time exploring the novel object (Figures S1I and S1J). Additionally, when the right chamber contained a novel object and the left chamber contained a familiar object, *Csnk2b*^+/-^ mice, like WT mice, exhibited a preference for exploring the novel object on the right side (Figures S1K and S1L).

We further examined the repetitive and stereotyped behaviors in *Csnk2b*^+/-^ mice through marble burying test and self-grooming recording. The results of the marble burying test showed no significant difference in the number of marbles buried between WT and *Csnk2b*^+/-^ mice within the same time period (Figures 1J and 1K). The self-grooming test also revealed no significant difference in self-grooming time between the two groups of mice (Figure 1L). Additionally, the open field test, which was used to assess anxiety levels in mice, showed that *Csnk2b*^+/-^ mice spent significantly less time in the central area compared to WT mice (Figures 1M and 1N), while their total locomotor distance was significantly increased (Figure 1O).

We assessed the spatial learning and memory abilities of *Csnk2b*^+/-^ mice using the Barnes maze test. During the first four days of training, there was no significant difference in the time required for *Csnk2b*^+/-^ mice to find the escape hole compared to WT mice (Figure 1P). On the fifth day of testing, the time taken by both groups of mice to locate the escape hole also showed no significant difference (Figure 1P). However, on the twelfth day of testing, the results indicated that *Csnk2b*^+/-^ mice required significantly more time to find the escape hole compared to WT mice (Figure 1P).

To investigate whether *Csnk2b*^+/-^ mice exhibit epileptic symptoms analogous to those observed in clinical patients, we continuously monitored their behavioral manifestations over a 24-hour period while simultaneously performing Electrocorticography (ECoG) recordings. A subset of 5- to 6-month-old *Csnk2b*^+/-^ mice (7 out of 10) displayed overt behavioral features of epileptic seizures, including wild running, myoclonic seizures, and generalized tonic-clonic seizures (Supplementary Video 1), accompanied by prominent epileptic discharges (Figures 1Q and 1R). In contrast, no spontaneous epileptic seizures were detected in WT mice (Figures 1Q and 1R). The frequency of spontaneous seizures in *Csnk2b*^+/-^ mice was 2.30±2.13 times/day, with a total daily seizure duration of 2.21±2.17 minutes (Figures 1S and 1T).

*Csnk2b* deficiency impairs the development of specific cortical neurons and neural network coordination To elucidate which specific neurons are affected in *Csnk2b*^+/-^ mice leading to the reduced cortical thickness, we stained neurons in various cortical layers. We first confirmed the significant reduction of *Csnk2b* expression in the cortex of *Csnk2b*^+/-^ mice through fluorescence staining (Figures 2A and 2B). Subsequently, using Cux1 staining, we found that the total number of neurons in layers 2 and 3 of the cortex was significantly reduced in *Csnk2b*^+/-^ mice compared to WT mice (Figures 2C and 2D). Counting neurons in layer 5 revealed a trend toward reduced numbers in *Csnk2b*^+/-^ mice, although this result was not statistically significant (Figures 2E and 2F). Finally, we observed a significant reduction in the number of neurons in layer 6 of the cortex in *Csnk2b*^+/-^ mice compared to WT mice (Figures 2G and 2H).

**Figure 2.**
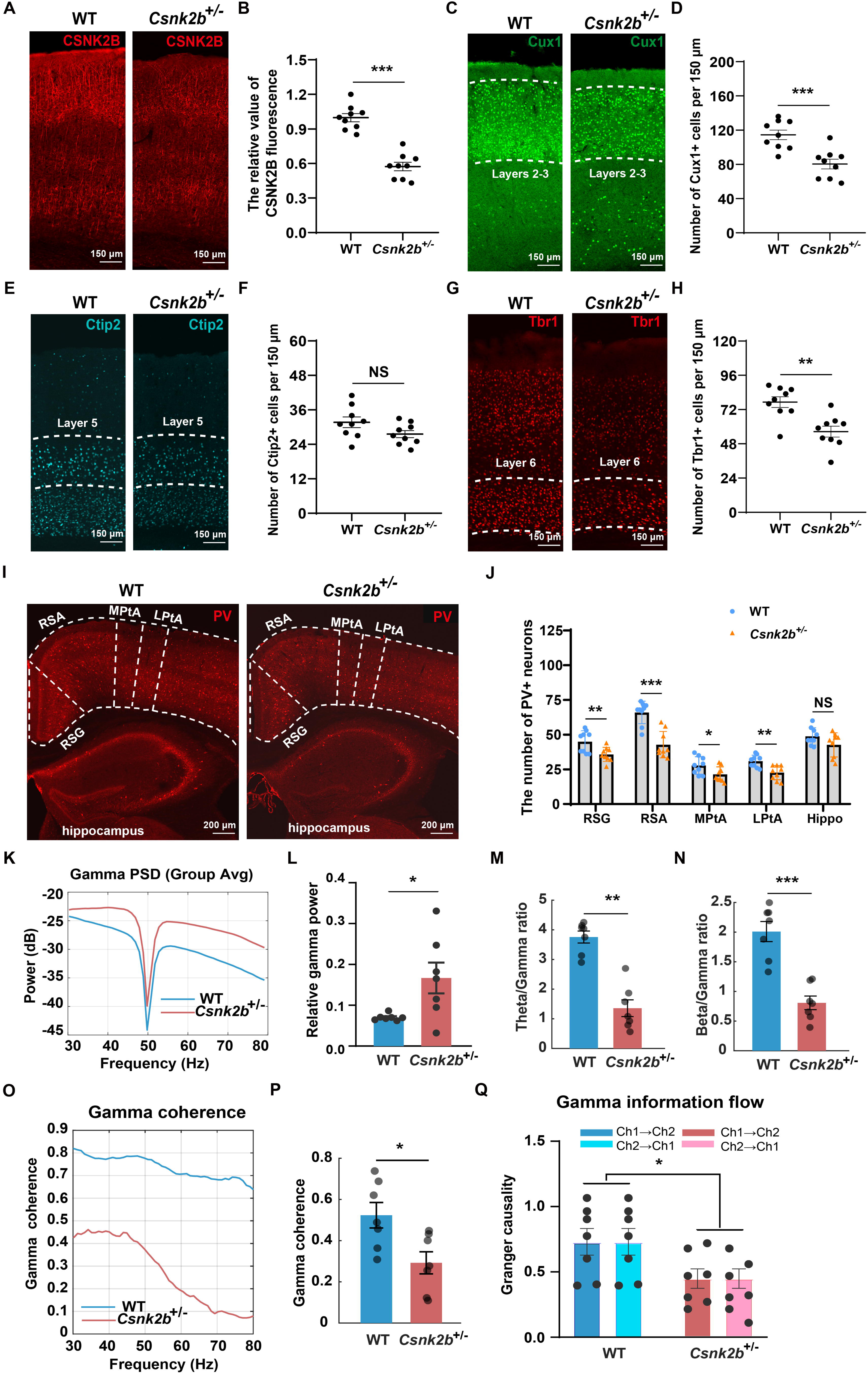
*Csnk2b* deficiency affects the development of specific cortical neurons and neural network communication. (A) Fluorescence staining of CSNK2B in the cortex of WT and *Csnk2b*^+/-^ mice. (B) Quantification of CSNK2B fluorescence intensity. (C) Staining of neurons in layers 2 and 3 of the cortex in WT and *Csnk2b*^+/-^ mice. (D) Quantification of neuronal numbers in layers 2 and 3 of the cortex. (E) Staining of neurons in layer 5 of the cortex in WT and *Csnk2b*^+/-^ mice. (F) Quantification of neuronal numbers in layer 5 of the cortex. (G) Staining of neurons in layer 6 of the cortex in WT and *Csnk2b*^+/-^ mice. (H) Quantification of neuronal numbers in layer 6 of the cortex. (I) Staining of PV-positive neurons in the cortex and hippocampus of WT and *Csnk2b*^+/-^ mice. (J) Quantification of PV-positive neurons in the cortex and hippocampus. (K) Group-averaged power spectral density (PSD) of ECoG signals recorded from the prefrontal cortex of WT and *Csnk2b*^+/-^ mice. (L) Quantification of gamma-band (30–80 Hz) relative power. (M) Theta/gamma ratio in WT and *Csnk2b*^+/-^ mice. (N) Beta/gamma ratio in WT and *Csnk2b*^+/-^ mice. (O) Frequency-dependent magnitude-squared coherence between bilateral PFC ECoG channels (Ch1 and Ch2) in the gamma band . (P) Mean gamma-band coherence values for WT and *Csnk2b*^+/-^ mice. (Q) Strength of bidirectional gamma-band information flow (assessed via Granger causality) between bilateral PFC channels (Ch1→Ch2 and Ch2→Ch1).Data are expressed as mean±SD, n = 9 slices from 3 mice (B, D, F, H, J), n = 7 mice for each group (L, M, N, P, Q). Statistical analysis was performed using two-tailed Student’s t-test. \**P <* 0.05, \*\**P <* 0.01, \*\*\**P <* 0.001, NS (not significant).

To further explore whether *Csnk2b* affects the development of specific inhibitory neurons, we analyzed its spatiotemporal expression patterns. We first utilized the ENCODE (the Encyclopedia of DNA Elements) data available on the NCBI website (https://www.ncbi.nlm.nih.gov/) and found that *Csnk2b* is expressed in both embryonic and adult mouse brains, with higher expression levels at embryonic day 14 (E14) (Figure S2A). Then, by consulting single-cell sequencing data from the Allen Brain Map database (https://portal.brain-map.org/), we discovered that *Csnk2b* is expressed in various inhibitory neurons, such as PV- and SST-positive neurons (Figure S2B). Staining for these two types of inhibitory neurons showed a significant reduction in PV-positive neurons in the cortex of *Csnk2b*^+/-^ mice, while no significant changes were observed in the hippocampus (Figures 2I and 2J). Additionally, compared to WT mice, the number of SST-positive neurons in both the cortical and hippocampal regions of *Csnk2b*^+/-^ mice remained unchanged (Figures S2C and S2D).

Since PV-positive neurons are pivotal for generating gamma oscillations and maintaining excitatory/inhibitory (E/I) balance, we reasoned that their loss would be reflected in aberrant network activity. Cortical ECoG recordings pointed to network-level hyperexcitability in *Csnk2b*^+/-^ mice. The E/I balance was fundamentally altered, as evidenced by a significant elevation in gamma-band (30-80 Hz) relative power (Figures 2K and 2L). This disinhibited state was further confirmed by a significant reduction in both the theta/gamma ratio and the beta/gamma ratio (Figures 2M and 2N).

We next asked whether this local disinhibition impaired large-scale network coordination. We found that long-range gamma synchrony, measured by magnitude-squared coherence between bilateral prefrontal sites, was severely compromised in *Csnk2b*^+/-^ mice (Figures 2O and 2P). To probe the functional impact of this desynchronization, we assessed the efficiency of directional information transfer using Granger causality. In both groups, the net information flow between channels was balanced (i.e., the difference between Channel 1→2 and Channel 2→1 was near zero). However, the analysis revealed a significant attenuation in the overall strength of this bidirectional gamma-band information flow in mutants (Figure 2Q).

### AAV-PHP.eB-mediated replacement therapy improves growth, cortical and hippocampal structure, and survival rate in *Csnk2b*^+/-^ mice

After observing the ASD-related behaviors and cellular phenotypes in *Csnk2b*^+/-^ mice, we investigated whether AAV-mediated gene therapy could ameliorate these deficits. Given that ASD is an early-onset neurodevelopmental disorder, we initiated treatment in *Csnk2b*^+/-^ mice at postnatal day 3. We selected AAV-PHP.eB, a serotype known for its efficient transduction across the blood-brain barrier. Additionally, we compared the therapeutic effects of the neuron-specific hsyn promoter and the non-cell-specific CAG promoter. The human *CSNK2B*-expressing AAV was delivered to the mouse brain via retro-orbital injection (Figure 3A).

**Figure 3.**
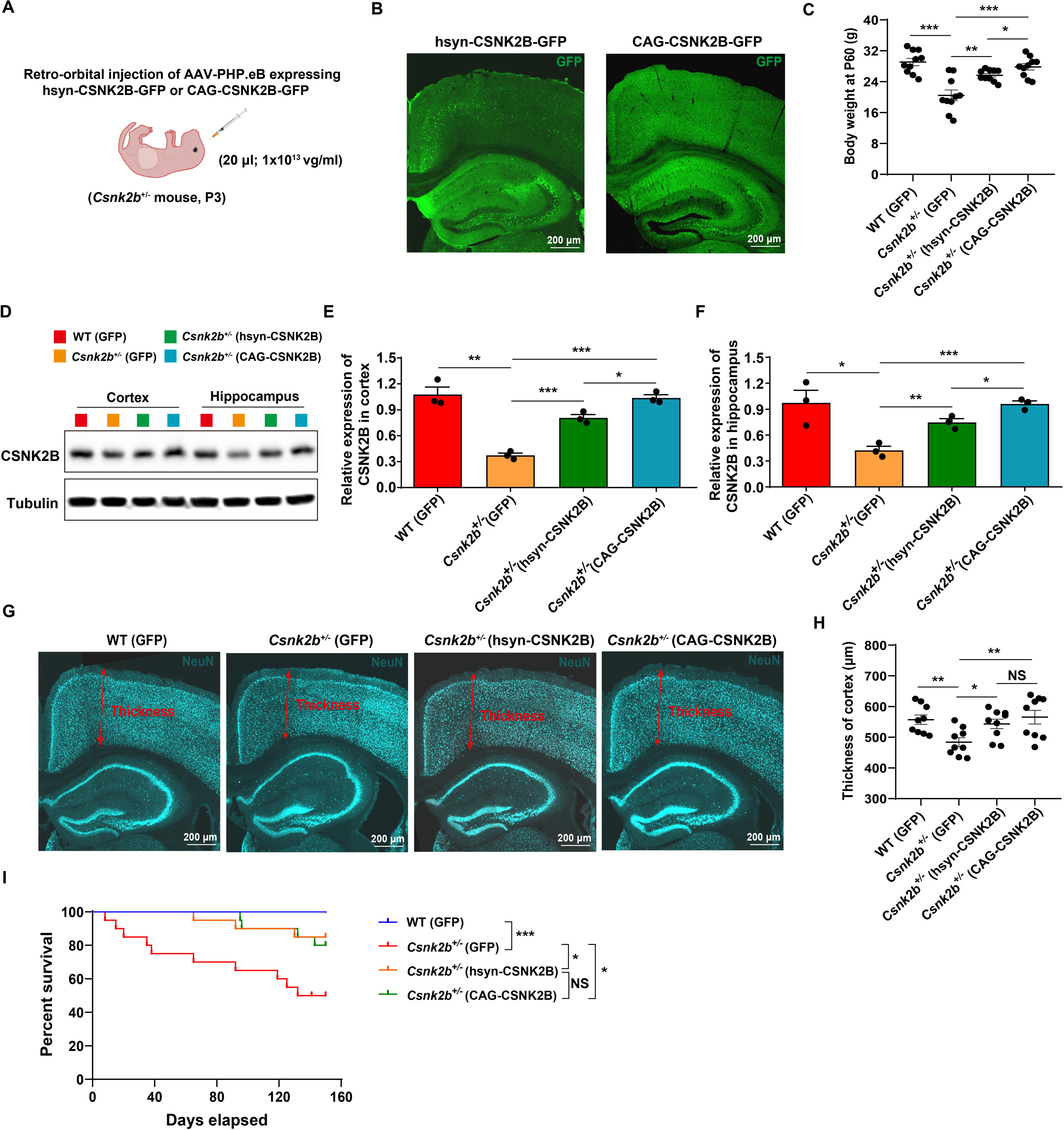
AAV-PHP.eB-mediated replacement therapy improves growth, cortical and hippocampal structure, and survival rate in *Csnk2b*^+/-^ mice. (A) Schematic illustration of AAV-PHP.eB-mediated replacement therapy. (B) Cellular infection in the cortex and hippocampus of *Csnk2b*^+/-^ mice following injection of AAVs carrying different promoters. (C) Body weight measurements of *Csnk2b*^+/-^ mice at 2 months of age following gene therapy (n = 10 mice for each group). (D) Detection of CSNK2B protein expression in the cortex and hippocampus of *Csnk2b*^+/-^ mice after injection of AAVs carrying different promoters. (E) Quantitative analysis of CSNK2B protein expression in the cortex (n = 3 mice for each group). (F) Quantitative analysis of CSNK2B protein expression in the hippocampus (n = 3 mice for each group). (G) Immunofluorescence staining of the cortex and hippocampus in *Csnk2b*^+/-^ mice after gene therapy. (H) Quantitative analysis of cortical thickness (n = 9 slices from 3 mice). (I) Survival analysis of *Csnk2b*^+/-^ mice up to 160 days following gene therapy (n = 20 mice for each group). Data are expressed as mean±SD. Statistical analysis was performed using one way ANOVA test (C, E, F, H) and log rank (Mantel-Cox) test (I) , with \**P <* 0.05, \*\**P <* 0.01, \*\*\**P <* 0.001 indicating significant differences.

One month after AAV injection, we found that AAV-CAG-CSNK2B infected a greater number of cells in the cortex and hippocampus compared to AAV-hsyn-CSNK2B (Figure 3B). Subsequent weight measurements showed that both AAV-hsyn-CSNK2B and AAV-CAG-CSNK2B injections significantly increased the body weight of *Csnk2b*^+/-^ mice, with AAV-CAG-CSNK2B having a pronounced more effect (Figure 3C). Quantification of protein expression revealed that both AAVs significantly increased CSNK2B protein levels in the cortex and hippocampus of *Csnk2b*^+/-^ mice, with AAV-CAG-CSNK2B yielding higher expression levels (Figures 3D-3F).

Fluorescence staining of the cortex and hippocampus indicated that both AAV treatments significantly increased cortical thickness in *Csnk2b*^+/-^ mice, although no significant difference was observed between the two treatments (Figures 3G and 3H).

Moreover, injection of AAV-hsyn-CSNK2B and AAV-CAG-CSNK2B improved the structure of the hippocampus and lateral ventricles in *Csnk2b*^+/-^ mice (Figure 3G). Finally, both AAV-hsyn-CSNK2B and AAV-CAG-CSNK2B treatments significantly enhanced the survival rate of *Csnk2b*^+/-^ mice, increasing it from 50% at 150 days postnatal to 80%, although no significant difference was detected between the two treatment groups (Figure 3I).

### AAV-PHP.eB-mediated replacement therapy rescues cortical neuron development in *Csnk2b*^+/-^ mice

Upon observing the recovery of cortical thickness in *Csnk2b*^+/-^ mice, we examined the number of neurons in each layer through immunofluorescence staining. We first confirmed that injection of AAV-hsyn-CSNK2B and AAV-CAG-CSNK2B significantly increased CSNK2B expression in the cortex, with AAV-CAG-CSNK2B yielding a higher expression level (Figures 4A and 4B). Subsequently, we found that the number of neurons in layers 2-3 of the cortex significantly increased in both treatment groups, although no significant difference was observed in the total number of neurons between the two groups (Figures 4C and 4D).

**Figure 4.**
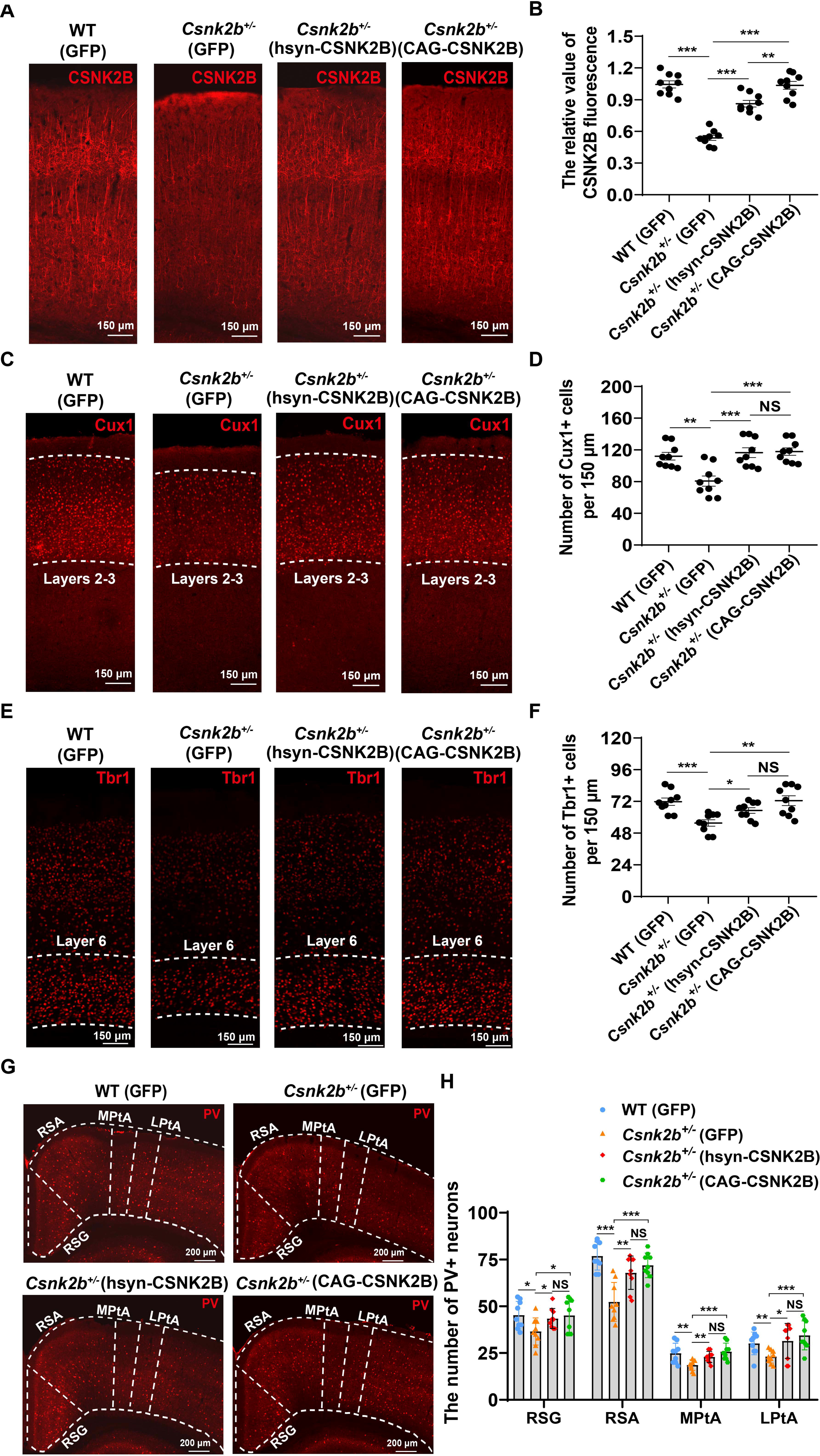
AAV-PHP.eB-mediated replacement therapy rescues cortical neuron development in *Csnk2b*^+/-^ Mice. (A) Immunofluorescence staining of CSNK2B in the cortex of *Csnk2b*^+/-^ mice after gene therapy. (B) Quantitative analysis of CSNK2B immunofluorescence intensity in the cortex. (C) Immunostaining of neurons in layers 2-3 of the cortex in *Csnk2b*^+/-^ mice after gene therapy. (D) Quantification of neuronal numbers in layers 2-3 of the cortex. (E) Immunostaining of neurons in layer 6 of the cortex in *Csnk2b*^+/-^ mice after gene therapy. (F) Quantification of neuronal numbers in layer 6 of the cortex. (G) Immunostaining of PV-positive neurons in the cortex of *Csnk2b*^+/-^ mice after gene therapy. (H) Quantification of PV-positive neurons in the cortex. Data are expressed as mean±SD, n = 9 slices from 3 mice. Statistical analysis was performed using one way ANOVA test, with \**P* < 0.05, \*\**P* < 0.01, \*\*\**P* < 0.001 indicating significant differences.

Additionally, after treatment with either AAV, the number of neurons in layer 6 of the cortex also significantly increased in *Csnk2b*^+/-^ mice, with no significant difference in treatment efficacy between the two AAVs (Figures 4E and 4F). Finally, we assessed the development of PV-positive neurons in the cortex of *Csnk2b*^+/-^ mice.

Compared to the control group, both AAV-hsyn-CSNK2B and AAV-CAG-CSNK2B treatments significantly increased the number of PV-positive neurons in the cortex, with no significant difference in the number of PV-positive neurons between the two treatment groups (Figures 4G and 4H).

### AAV-PHP.eB gene replacement normalizes EEG/ECoG biomarkers of E/I balance and large-scale network coordination

Because PV-interneuron loss in *Csnk2b*^+/-^ cortex predicted oscillatory disruption, we asked whether neonatal AAV-PHP.eB–CSNK2B would correct EEG-detectable network signatures that could serve as translational biomarkers. A two-way ANOVA on band-limited relative power revealed a significant Groups × Frequency Band interaction (F (8, 118) = 2.922, *p* = 0.005), indicating frequency-specific rescue. Post-hoc tests showed selective normalization of two prespecified endpoints: the pathological gamma elevation was reduced and the suppressed theta power was restored (Figures 5A and 5B). While gamma peak frequency and bandwidth exhibited only trends toward reversal (Figures 5C and 5D), composite ratios that index cortical E/I balance were robustly corrected—most notably, the theta/gamma ratio, which was reduced in *Csnk2b*^+/-^ mice, returned to WT levels after treatment (Figure 5E). The beta/gamma ratio showed a concordant, though nonsignificant, tendency toward normalization (Figure 5F). Together these data nominate theta and gamma band power—and their ratios—as sensitive EEG biomarkers of target engagement and physiological efficacy for CSNK2B-linked neurodevelopmental disorders.

**Figure 5.**
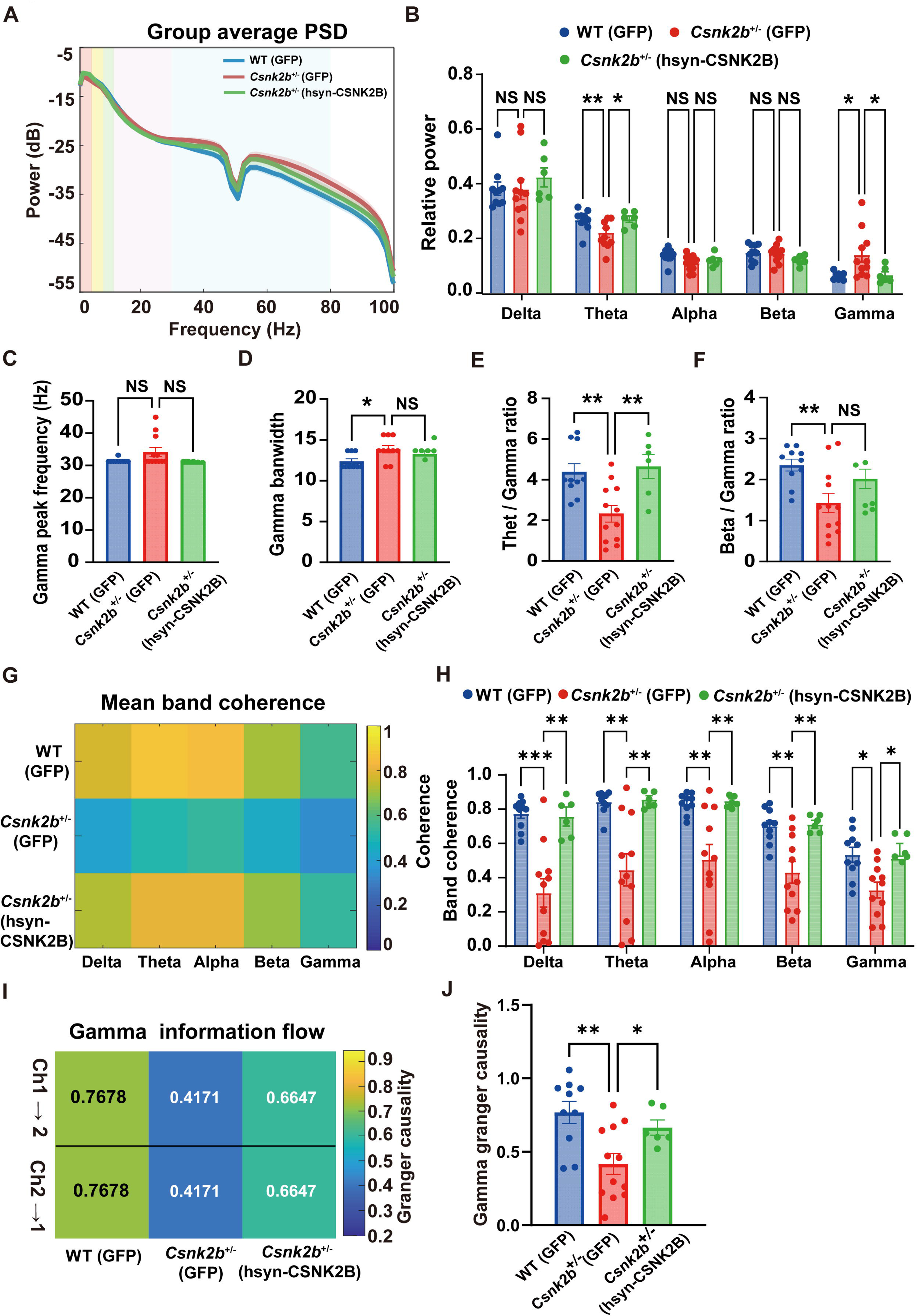
AAV-PHP.eB-mediated replacement therapy restores cortical E/I balance and global network synchronization. (A) Group-averaged power spectral density (PSD) of ECoG signals recorded from the prefrontal cortex of mice. (B) Quantification of relative power across different frequency bands (Delta, Theta, Alpha, Beta, Gamma) in the three groups. (C) Quantification of gamma oscillation peak frequency. (D) Quantification of gamma bandwidth. (E) Quantification of the theta/gamma ratio in the three groups. The theta/gamma ratio is a critical marker of E/I balance. (F) Quantification of the beta/gamma ratio in the three groups. The beta/gamma ratio is an additional index of E/I balance. (G) Frequency-dependent magnitude-squared coherence between bilateral PFC ECoG channels (Ch1 and Ch2) across different frequency bands in the three groups. Coherence reflects inter-regional neural synchronization. (H) Quantification of mean band coherence across Delta, Theta, Alpha, Beta, and Gamma frequency bands in the three groups. (I) Strength of bidirectional gamma-band information flow (assessed via Granger causality) between bilateral PFC channels (Ch1→Ch2 and Ch2→Ch1) in the three groups. (J) Quantification of the absolute strength of bidirectional gamma-band information flow in the three groups. Data are expressed as mean ± SD, n = 6-12 mice for each group. Statistical analysis was performed using one way ANOVA test (C, D, E, F, J) and two-way ANOVA test (B, H). \**P <* 0.05, \*\**P <* 0.01, \*\*\**P <* 0.001, NS (not significant).

We next evaluated whether local circuit repair generalized to macroscopic communication. Two-way repeated-measures ANOVA of magnitude-squared coherence demonstrated a strong main effect of Group (F (2, 23) = 14.15, *p* < 0.0001) and a significant Groups × Frequency Band interaction (F (8, 92) = 3.513, *p* = 0.0014), consistent with band-specific network impairment and rescue. Gene therapy increased inter-regional synchrony and restored coherence to WT levels across key bands, including gamma (Figures 5G and 5H), positioning band-limited coherence as a second quantitative biomarker class.

Finally, to probe effective connectivity, we performed Granger causality focused on gamma (a PV-dependent rhythm). Although net directionality remained near zero in all groups (balanced bidirectional flow; Figure 5I), the absolute strength of bidirectional gamma-band influence—a measure of information transfer efficiency—was significantly reduced in *Csnk2b*^+/-^ mice and was restored by gene therapy to WT levels (Figure 5J). Notably, all analyses replicated in light-phase data, indicating state-independent robustness of these EEG endpoints (Figures S3A-S3J).

Collectively, these results define a compact, translatable EEG biomarker panel—(i) theta and gamma relative power, (ii) theta/gamma (± beta/gamma) ratios, (iii) band-limited coherence, and (iv) gamma-band Granger causality—that reports AAV target engagement, E/I rebalancing, and network-level rescue. These signatures offer objective efficacy readouts to guide dose optimization and early clinical decision-making in gene therapy for neurodevelopmental disorders.

### AAV-PHP.eB-mediated replacement therapy rescues autistic-like behavioral phenotypes and spontaneous epilepsy in *Csnk2b*^+/-^ mice

After the *Csnk2b*^+/-^ mice received treatment, we first assessed their social ability using the three-chamber test. During the sociability test, both groups of mice exhibited a preference for interacting with the stranger mouse rather than the empty cage (Figures 6A and 6B). Additionally, during the social novelty test, compared to the control group (*Csnk2b*^+/-^ mice injected with AAV-GFP), both treatment groups showed a significant social preference, spending significantly more time interacting with the stranger mouse than with the familiar mouse (Figures 6C and 6D).

**Figure 6.**
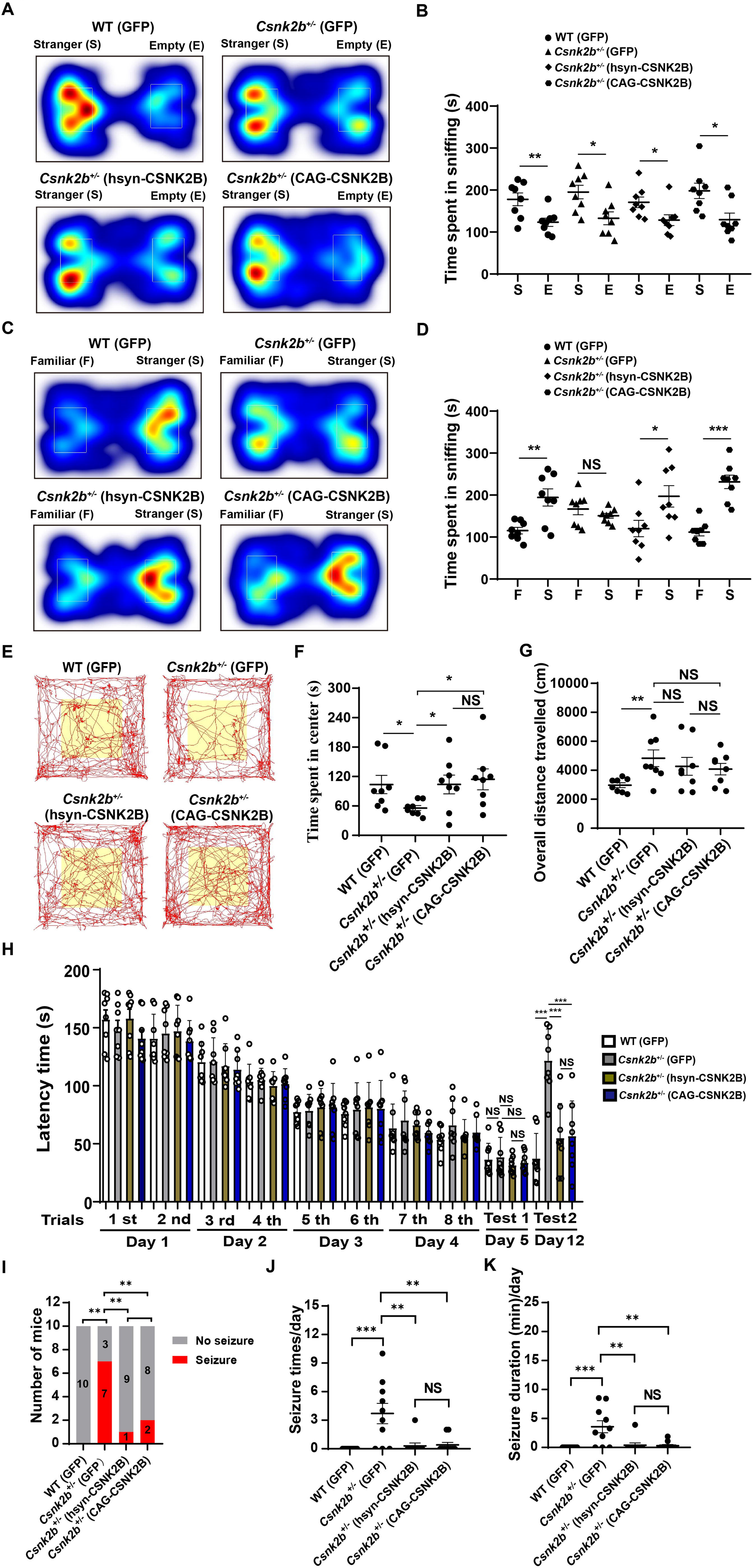
AAV-PHP.eB-mediated replacement therapy rescues autistic-like behavioral phenotypes and spontaneous epilepsy in *Csnk2b*^+/-^ mice. (A) Representative heatmaps of mouse trajectories during the sociability test. (B) Time spent interacting with the stranger mouse and empty cage during the sociability test. (C) Representative heatmaps of mouse trajectories during the social novelty test. (D) Time spent interacting with the stranger and familiar mice during the social novelty test. (E) Representative trajectories of mice in the open field test. (F) Time spent exploring the central area in the open field test for each group. (G) Total distance traveled in the open field test for each group. (H) Quantitative analysis of time taken to find the escape hole on day 12 in the Barnes maze test for each group. (I) Incidence of spontaneous seizures in *Csnk2b*^+/-^ mice following neonatal injection of AAV-hsyn-CSNK2B or AAV-CAG-CSNK2B. (G) Daily frequency of spontaneous seizures in *Csnk2b*^+/-^ mice after neonatal AAV-mediated replacement therapy. (K) Total daily duration of spontaneous seizures in *Csnk2b*^+/-^ mice after neonatal AAV-mediated replacement therapy Data are expressed as mean±SD, n = 8 mice for each group (B, D, F, G, H), n = 10 mice for each group (I, J, K). Statistical analysis was performed using two-tailed Student’s t-test (B, D), one way ANOVA test (F, G, H), Fisher’s exact test (I) and Dunn’s test (J, K), with \**P* < 0.05, \*\**P* < 0.01, \*\*\**P* < 0.001 indicating significant differences. NS (not significant).

Subsequently, we evaluated the anxiety levels of the treated mice using the open field test. After injection with AAV-hsyn-CSNK2B or AAV-CAG-CSNK2B, *Csnk2b*^+/-^ mice spent significantly more time exploring the central area of the open field, although no significant difference was observed between the two treatment groups (Figures 6E and 6F). However, compared to the control group, the total distance traveled in the open field by both treatment groups did not show a significant change, and no significant difference was detected between the two treatment groups in terms of locomotor activity (Figure 6G).

We then investigated whether the long-term spatial learning and memory abilities of *Csnk2b*^+/-^ mice were improved by using the Barnes maze test. We found that after injection of AAV-hsyn-CSNK2B or AAV-CAG-CSNK2B, the time required for *Csnk2b*^+/-^ mice to locate the escape hole in the test on Day 12 was significantly reduced (Figure 6H). However, no significant difference was observed in the time taken to find the escape hole between the two treatment groups (Figure 6H). In addition, during the test on Day 5, there was no significant difference in the time taken to locate the escape hole among the four groups of mice (Figure 6H), which indicates that the short-term spatial learning and memory abilities of *Csnk2b*^+/-^ mice were not impaired.

Finally, we evaluated the 24-hour seizure profiles of *Csnk2b*^+/-^ mice at 5-6 months of age following gene therapy administered at the neonatal stage (P3). The results demonstrated that injection of either AAV-hsyn-CSNK2B or AAV-CAG-CSNK2B significantly reduced the incidence of spontaneous seizures in *Csnk2b*^+/-^ mice (Figure 6I). The seizure incidence decreased from 70% (7 out of 10) in the control group to 10% (1 out of 10) in the AAV-hsyn-CSNK2B group and 20% (2 out of 10) in the AAV-CAG-CSNK2B group, with no significant difference between the two therapeutic groups (Figure 6I). Furthermore, relative to the control group, both therapeutic groups exhibited marked reductions in seizure frequency and total seizure duration (Figures 6J and 6K), whereas no significant differences were detected between the two AAV-treated groups (Figures 6J and 6K). Specifically, the daily seizure frequency of *Csnk2b*^+/-^ mice decreased from 3.70±3.21 times/day (control group) to 0.30±0.95 times/day (AAV-hsyn-CSNK2B group) and 0.40±0.84 times/day (AAV-CAG-CSNK2B group) (Figure 6J). Correspondingly, the total daily seizure duration was reduced from 3.57±3.43 minutes per day to 0.39±1.23 minutes per day and 0.30±0.59 minutes per day (Figure 6K).

## Discussion

In this study, we successfully generated a heterozygous knockout mouse model of *Csnk2b* and conducted a comprehensive investigation into its behavioral, neurodevelopmental, and epileptic phenotypes. Furthermore, we evaluated the therapeutic potential of AAV-mediated gene replacement. Our findings demonstrate that *Csnk2b* haploinsufficiency in mice leads to autism-like behaviors, epileptic phenotypes, growth retardation, structural brain abnormalities, specific cortical neuronal deficits, and network desynchronization. Notably, early postnatal administration of AAV-PHP.eB-mediated gene therapy effectively ameliorated these deficits. These results provide critical preclinical evidence for elucidating the pathological mechanisms underlying *CSNK2B*-related neurodevelopmental disorders and support the development of targeted therapeutic strategies.

The core phenotypes observed in *Csnk2b*^+/-^ mice closely mirror the clinical manifestations of human disorders associated with *CSNK2B* mutations. At the behavioral level, these mice exhibited impaired social novelty recognition and heightened anxiety-like behaviors, which are consistent with the social deficits and emotional dysregulation commonly observed in individuals with ASD. Social novelty recognition is a complex cognitive process that depends on the coordinated function of brain regions such as the prefrontal cortex and basolateral amygdala, as well as specialized neuronal circuits, including the PV-positive interneuron network^18,19^. The significant reduction in cortical neurons across layers 2/3 and 6, as well as the specific loss of PV-positive interneurons observed in our study, may disrupt the integrity of these circuits, thereby contributing to the observed social behavioral impairments. Moreover, *Csnk2b*^+/-^ mice showed deficiencies in long-term spatial learning and memory, had epileptic seizures, and exhibited lower body weight. These characteristics are analogous to the intellectual disability, epilepsy, and growth retardation seen in patients with POBINDS. These findings suggest that *Csnk2b*^+/-^ mice represent a valuable model for studying diseases linked to *CSNK2B* mutations.

At the neurodevelopmental level, our findings revealed that *Csnk2b* haploinsufficiency led to a reduction in brain weight, cortical thinning, and structural abnormalities in the hippocampus. Previous research has shown that *Csnk2b* deficiency impairs the proliferation and differentiation of neural stem cells^11,15^, which may be the underlying cause of the observed defects in brain development. Beyond neural stem cells, CSNK2B also plays a role in modulating the proliferation of diverse cell types, such as hepatocytes and colorectal cells^20,21^. This suggests that *Csnk2b* can modulate the proliferation processes of multiple cell types, thereby leading to systemic growth retardation in *Csnk2b*^+/-^ mice.

Importantly, our study revealed that *Csnk2b* deficiency differentially impacts the development of specific cortical neuronal subtypes. We observed a significant reduction in neurons within the superficial layers (layers 2/3) and deep layers (layer 6) of the cortex, as well as a specific loss of PV-positive inhibitory interneurons. This selective vulnerability may be attributed to the unique signaling dependencies of these neuronal populations. For instance, the mTOR and Wnt/β-catenin pathways, both of which are regulated by *CSNK2B*, are known to play critical roles in the development and function of PV-positive interneurons^20,22–25^. The reduction of PV-positive inhibitory interneurons is particularly noteworthy, as these cells are crucial for maintaining the E/I balance in cortical circuits^26^. Our data also confirm the cortical E/I imbalance in *Csnk2b*-deficient mice, characterized by increased gamma power and a decreased theta/gamma ratio. PV-positive neurons are crucial for fast inhibition and gamma rhythm generation^27,28^. This microcircuit dysfunction not only leads to local hyperexcitability but also impairs global brain communication. This impairment not only constitutes a potential mechanism contributing to autism-like behaviors but also provides a plausible explanation for the increased seizure susceptibility observed in the model^29–32^.

Our findings demonstrate that AAV-PHP.eB-mediated replacement therapy can effectively reverse multiple phenotypes in *Csnk2b*^+/-^ mice. From a therapeutic strategy standpoint, the timing of intervention during the neonatal period is critical. At this stage, the brain is still undergoing rapid development, with neural stem cells actively proliferating and differentiating^33,34^. The introduction of exogenous genes at this time can timely rescue neuronal developmental abnormalities. Regarding serotype selection, we opted for the AAV-PHP.eB variant, which was engineered from . Its enhanced capability to efficiently cross the blood-brain barrier enables whole-brain delivery via retro-orbital injection, thereby circumventing the potential damage associated with invasive intracranial injections in neonatal mice. Compared to the AAV9 serotype, AAV-PHP.eB exhibits superior transduction efficiency in the central nervous system and poses a lower risk of hepatotoxicity^36,37^. However, AAV-PHP.eB also has certain limitations. For example, its high efficiency is strain-dependent in mice^36,38^, and in non-human primates (such as macaques), its transduction efficiency does not exhibit a significant advantage^39,40^. These factors may restrict its broad applicability across different animal models or human populations.

We compared the therapeutic efficacy of a neuron-specific promoter (hsyn) with that of a broad-spectrum strong promoter (CAG). Although AAV-CAG-CSNK2B showed higher infection rates and protein expression levels than AAV-hsyn-CSNK2B, there were no significant differences between the two treatments in cortical neuron development, behavioral phenotypes, epileptic symptoms, and survival rates. Possible explanations include: (1) CSNK2B protein or its regulatory functions may have redundancy or a threshold effect, where reaching a critical expression level can effectively support neuronal development and function; (2) The high expression of AAV-hsyn-CSNK2B in neurons may be sufficient to drive necessary biological effects, while the additional infection of non-neuronal cells by AAV-CAG-CSNK2B may contribute little to core phenotype improvements; (3) Both vectors may achieve similar functional *CSNK2B* expression levels in key pathological cell types (especially PV-positive neurons), sufficient to correct defects. This finding suggests that both neuron-specific and non-specific promoters may be effective in future therapies, offering flexibility in vector design.

Certainly, this study has several limitations. Firstly, our focus was primarily on the macroscopic structures of the cortex and hippocampus, as well as specific neuronal types. Future research should delve deeper into the specific mechanisms by which *Csnk2b* deficiency impacts particular neural circuits, synaptic plasticity, and downstream signaling pathways. Secondly, the model only simulates *CSNK2B* heterozygous deletion, whereas human patients exhibit a variety of mutations, including splice mutations and missense mutations, which may have differing functional impacts. Lastly, the intervention in this study was conducted at a very early postnatal stage (P3), simulating a preventive or extremely early treatment scenario. The therapeutic efficacy for individuals with already apparent symptoms remains to be further investigated. Subsequent studies should aim to elucidate more refined molecular mechanisms, explore the efficacy of treatment within a broader therapeutic time window, and advance translational research toward clinical applications.

## Materials and methods

### Animal

All animal experiments were conducted in strict accordance with the guidelines approved by the Animal Care and Ethical Committee of Songjiang Hospital Affiliated to Shanghai Jiao Tong University School of Medicine (Approval No. ACE-004-2025). Mice were housed in a specific pathogen-free (SPF) animal facility, maintained under a 12-hour light/12-hour dark cycle (lights on from 7:00 AM to 7:00 PM) and kept at a controlled temperature of 22–25°C. The genotypes of *Csnk2b*^+/-^ mice with a C57BL/6J background were confirmed by PCR using the following primers: forward primer 5’-TGTTGTTTGAGAGCAGATTTCGAC-3’ and reverse primer 5’-CAGTTTGGGTTATGAATGCTTGGG-3’. Behavioral tests were performed during the light phase, with age-matched (2-3 months old) and genotyped mice being tested in a blinded manner. To acclimate the mice to the experimental environment, they were handled for 3 days prior to behavioral testing and were allowed to adapt to the testing room for 30 minutes before the tests.

### Three-chamber test

The three-chamber social test assesses social preference and social novelty preference in animals. Mice were acclimated to a rectangular chamber divided into three sections (left, middle, and right) for 10-15 minutes, 1-3 days before the experiment. For the social preference test, a cage containing a stranger mouse (S1) was placed in the left section, and an empty cage in the right section. The experimental mouse was introduced into the middle section, and the time spent sniffing the mouse and cage in the left and right sections was recorded over 10 minutes. Normal animals typically prefer S1. For the social novelty test, the empty cage was replaced with a cage containing a novel stranger mouse (S2), and social interaction time was recorded over another 10 minutes. Normal animals generally prefer S2.

### Novel object recognition test

This test was performed in a box (40 cm width × 60 cm length) with three equal-sized chambers. The experiment process consisted of three parts, namely habituation, object cognition, and novel object recognition. During the first part, an empty cage was put into each side chamber, followed by letting the test mouse explore the box for 10 min. For the second part, a green cone block (object 1) was placed into the left empty cage, and then the test mouse was allowed to explore freely for 10 min. Lastly, after a pink cone block (new object) was put into the right empty cage, the test mouse was given 10 min to explore. The time spent by mice exploring objects was analyzed using Noldus EthoVision XT 11.5 software.

### Marble burying test

This test was performed in a clean box (40 cm width × 40 cm length) with a 7 cm thick layer of bedding, and 25 glass marbles were placed on the bedding surface. The test mouse was free to explore the glass marbles for 10 minutes. The glass marble that was covered off at least 50% was counted.

### Self-grooming recording

We put the test mouse into a clean cage without food or water, and recorded its activity in the cage for 30 min using a high-definition camera. Counting grooming behaviors included wiping faces, scratching heads and ears, and grooming the entire body. The grooming time was manually counted by using a stopwatch.

### Open field test

The test mouse was placed in a clean box (40 cm width × 40 cm length) for 10 min of habituation. After that, the activity of mouse was recorded by a camera for 10 min. The time spent in central area (20 cm width × 20 cm length) of the box was analyzed by Noldus EthovisonXT 11.5 software.

### Barnes maze

The Barnes maze comprises a 2-meter-diameter round table with 40 evenly spaced small holes around its perimeter. A shaded escape box is placed beneath one of these holes. A bright light is positioned above the table to prompt the test mouse to find the escape box. The experiment includes a 4-day training session and tests on days 5 and 12. The mice were trained twice each day for the first four days. During training, the mouse is placed in the table center and released from a black box to explore freely for 3 minutes. The time to find the escape box is recorded, followed by a 1-minute rest in the box. If the mouse fails to find the box within 3 minutes, it is still placed in the box for 1 minute. After each training trial, the table is cleaned and rotated to change the position of escape hole. On test days 5 and 12, the time to find the escape box is recorded.

### Immunofluorescence assays

Mice were first anesthetized and perfused transcardially with cold PBS, followed by perfusion with 4% cold paraformaldehyde (PFA). The brains were then extracted and post-fixed overnight at 4°C in 4% PFA. After fixation, the brains were washed with PBS and dehydrated sequentially in 15% sucrose for 24 hours and 30% sucrose for 48 hours. The dehydrated brains were embedded in optimal cutting temperature (OCT) compound (SAKURA, 4583) and frozen in a cryostat chamber (Leica, CM1950) at -20°C. The frozen brains were sectioned at a thickness of 40 µm at -25°C. For staining, the brain sections were first blocked with PBS containing 5% BSA and 0.3% Triton X-100 for 1.5 hours at room temperature. They were then incubated overnight at 4°C with the following primary antibodies: Csnk2b (1:500, Abcam, ab76025), Cux1 (1:500, Abcam, ab307821), Ctip2 (1:500, Abcam, ab18465), Tbr1 (1:500, Abcam, ab183032), PV (1:1000, Abcam, ab181086), NeuN (1:500, Millipore, ABN78), SST (1:250, Santa Cruz, sc-55565). After three washes with PBS, the sections were incubated with fluorescence-labeled secondary antibodies for 1 hour at room temperature. All antibodies were diluted in PBS containing 3% BSA and 0.1% Triton X-100. Finally, the brain sections were mounted on microscope slides using Fluoromount-G (Southern Biotechnology, 0100-01).

### Western Blot

To extract proteins, brain tissue was homogenized in lysis buffer (Beyotime, P0013) containing 1% protease inhibitor cocktail (Beyotime, P0015). Protein concentrations were determined using a BCA Protein Assay Kit (Beyotime, P0010S). Proteins were separated on 12% BeyoGel™ Plus PAGE gels (Beyotime, P0458) and transferred onto PVDF membranes (Millipore, IPVH00010). The membranes were blocked with 5% milk for 1 hour at room temperature, followed by overnight incubation with primary antibodies at 4°C. The information of the primary antibody is as follows: Csnk2b (1:1000, Abcam, ab76025), β-Tubulin (1:3000, Abcam, ab6046). After three washes with PBST, the membranes were incubated with secondary antibodies for 1 hour at room temperature. Protein bands were visualized using a chemiluminescent substrate (Thermo Fisher Scientific, 32106).

### Real-Time *Quantitative* PCR

Total RNA was extracted from mouse tissue using the RNAsimple Total RNA Kit (TIANGEN, DP419). cDNA synthesis was performed using the HiScript III 1st Strand cDNA Synthesis Kit (Vazyme, R312-01). Gene expression levels were quantified using ChamQ Universal SYBR qPCR Master Mix (Vazyme, Q711) according to the protocol of manufacturer. The housekeeping gene Gapdh was used as an internal control. The cycling conditions were as follows: 30 seconds at 95°C, followed by 40 cycles of 95°C for 10 seconds and 60°C for 30 seconds. Primers for *Csnk2b* were as follows: forward primer 5’-GCAGGTGCCTCACTATCGAC-3’ and reverse primer 5’-AAGCATCTCAG CTGCCTGTT-3’. Primers for *Gapdh* were as follows: forward primer 5’-GTGAAGGT CGGTGTGAACGG-3’ and reverse primer 5’-CGCTCCTGGAAGATGGTGAT-3’.

### AAV Injection

The AAV-PHP.eB-hSyn-CSNK2B-EGFP and AAV-PHP.eB-CAG-CSNK2B-EGFP were obtained from PackGene Biotech for retro-orbital injection. A total volume of 20 µL per pup (viral titer: 1 × 10¹³ vg/mL) was administered to neonatal mice on postnatal day 3. Based on published protocols^41^, the procedure for retro-orbital injection in newborn mouse pups is as follows:

First, draw the appropriate volume of viral solution into a 31-gauge insulin syringe. Temporarily transfer the dam to a clean holding cage, leaving the pups in the original home cage. Anesthetize each pup by placing it on ice for approximately 3 minutes, until the skin turns slightly bluish, indicating sufficient anesthesia. Next, place the pup in left lateral recumbency on a clean paper towel with its head facing to the right. Gently restrain the head of pup using the thumb and forefinger of the non-dominant hand, taking care not to compress the trachea or obstruct venous flow. Locate the right eye region (typically at the 3 o’clock position of the eyeball), and insert the needle at a 30-45° angle with the bevel facing down, to a depth of approximately one-third of the needle length (∼3 mm). Slowly and steadily inject the viral solution. After injection, leave the needle in place for 3-5 seconds before gently withdrawing it. Place the pup on a 37°C warming pad to recover body temperature until it regains mobility and normal pink coloration. Finally, gently rub the pup with nesting material from the home cage to remove any foreign scent before returning it to the dam, thereby preventing maternal rejection due to unfamiliar odor.

### ECoG recording

Mice were anesthetized via intraperitoneal injection and then fixed on a stereotaxic instrument. The hair on the top of the head was shaved, and the area was disinfected with povidone-iodine. The scalp was incised and separated to expose the skull. Four holes were drilled vertically into the skull using a cranial drill, followed by screwing in screws. Subsequently, the electrode wires of a 4-channel mouse electrode (Bio-Signal Technologies, 2623) were wrapped around the screws. Two recording electrodes were strategically implanted bilaterally over the prefrontal cortices (PFC) at coordinates: Left PFC (AP: +2.0 mm, ML: -1.5 mm); Right PFC (AP: +2.0 mm, ML: +1.5 mm). A cerebellar reference electrode (AP: -6.0 mm, ML: 0.0 mm) was placed to minimize cortical signal contamination, with a ground wire secured posterior to lambda. Dental cement was used to fix the screws and the mouse electrode. After the cement solidified, the mice were placed back on a heating pad for recovery. ECoG signals from seizure mice were acquired using a Medusa electrophysiological recording system (Bio-Signal Technologies), the analysis of seizure frequency and total seizure duration was performed using Sirenia Seizure software.

### Data acquisition and preprocessing

After 24-hour cable habituation, continuous ECoG signals were acquired across a full 24-hour circadian cycle using a Pinnacle Technology setup, sampled at 1000 Hz with online band-pass filtering (0.5-500 Hz) and 50/60 Hz notch filtering. Synchronized video was recorded throughout for behavioral correlation. Data from all groups were processed through a standardized MATLAB pipeline: raw EDF signals were downsampled to 250 Hz and segmented into 60-minute epochs. The first two ECoG channels from the 9th nocturnal segment (hours 8-9) were selected for core analysis, followed by multi-notch filtering (50 Hz harmonics elimination), 4th-order Butterworth band-pass filtering (0.5-100 Hz), and z-score normalization to ensure signal quality.

### Spectral power analysis

Power spectral density was computed using Welch’s method (256-point window, 128-point overlap, 256-point FFT) across five key frequency bands: δ (1-4 Hz), θ (4-8 Hz), α (8-13 Hz), β (13-30 Hz), and γ (30-80 Hz). Both absolute power and relative power (normalized to total power in 1-100 Hz range) were analyzed for each band, providing fundamental data for subsequent E/I imbalance metric calculations.

### Coherence Analysis

Magnitude-squared coherence was employed to quantify the linear dependence and frequency-specific synchronization between the two ECoG channels. The coherence estimate, *C_XY_* (*f*), between signals *x(t)* and *y(t)* at frequency f was calculated using the following formula:

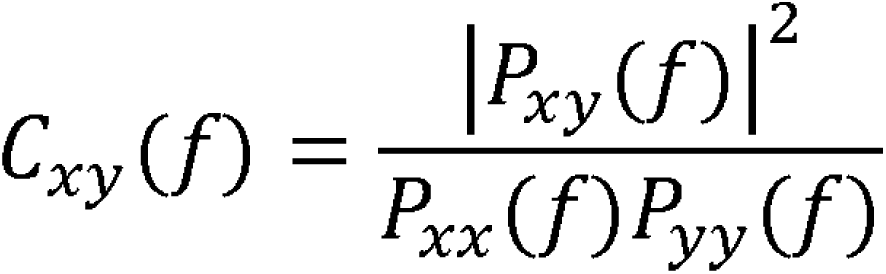

where *P_XX_* (*f*) and *P_YY_*(*f*) are the power spectral densities of *x* and *y*, and *P_XY_* (*f*) is the cross-power spectral density. Calculations were based on Welch’s periodogram method (256-point Hamming window, 50% overlap), yielding coherence values ranging from 0 (no connectivity) to 1 (perfect synchronization). The mean coherence within each pre-defined frequency band was subsequently computed for group-level statistical comparisons.

### Granger causality analysis

Directional information flow was investigated using Granger causality analysis. We fitted a bivariate vector autoregressive (VAR) model with an order of 8 to the preprocessed signals. Conceptually, a signal X is said to Granger-cause a signal Y if the prediction of Y is improved by incorporating past values of X. By comparing the predictive power of models that included or excluded the historical information from one channel to another, we quantified the unidirectional causal influence (Granger causality) in both directions (GCX→Y and GCY→X). The net causal flow was subsequently defined as the difference between these two directional influences (GCX→Y − GCY→X), providing a measure of the dominant direction of information exchange.

### Statistical analysis

All data were analyzed using GraphPad Prism 8 software and are presented as mean ± standard deviation (SD), with the sample size (n) specified in each figure legend. For comparisons between two independent groups, two-tailed Student’s t-test, Fisher’s exact test, and Mann-Whitney U test were applied. one-way ANOVA and Dunn’s test were used for comparisons among data of more than two groups. A p-value of less than 0.05 was considered statistically significant and is indicated as follows: \**p* < 0.05, \*\**p* < 0.01, \*\*\**p* < 0.001, and NS denotes not significant.

### Data and code availability

The raw data required to reproduce these findings will be available at the corresponding author’s request.

## Acknowledgments

This work was supported by the following grants: National Science and Technology Major Project (2025ZD0214700), National Natural Science Foundation of China (#82301334 CD, #82430046 ZQ), National Science and Technology Major Project (2025ZD0214700), and Project of Medical Technology Research and Transformation supported by Shanghai Municipal Health Commission (2024ZZ1007).

## Author contributions

The study was designed by Z.Q., A.D., and C.D.. C.D. conducted behavioral tests, immunohistochemistry, and virus injections. Y.Y., Y.Z., and Y.H. were responsible for the mouse breeding and genotyping. X.W. assisted in conducting ECoG recordings and the associated data analysis. The manuscript was written by C.D. and Z.Q..

