## Supplemental information for "AAV-mediated *CSNK2B* gene replacement rescues ASD-relevant phenotypes and establishes EEG biomarkers for translation in *Csnk2b* haploinsufficient mice"

Supplementary Figure1

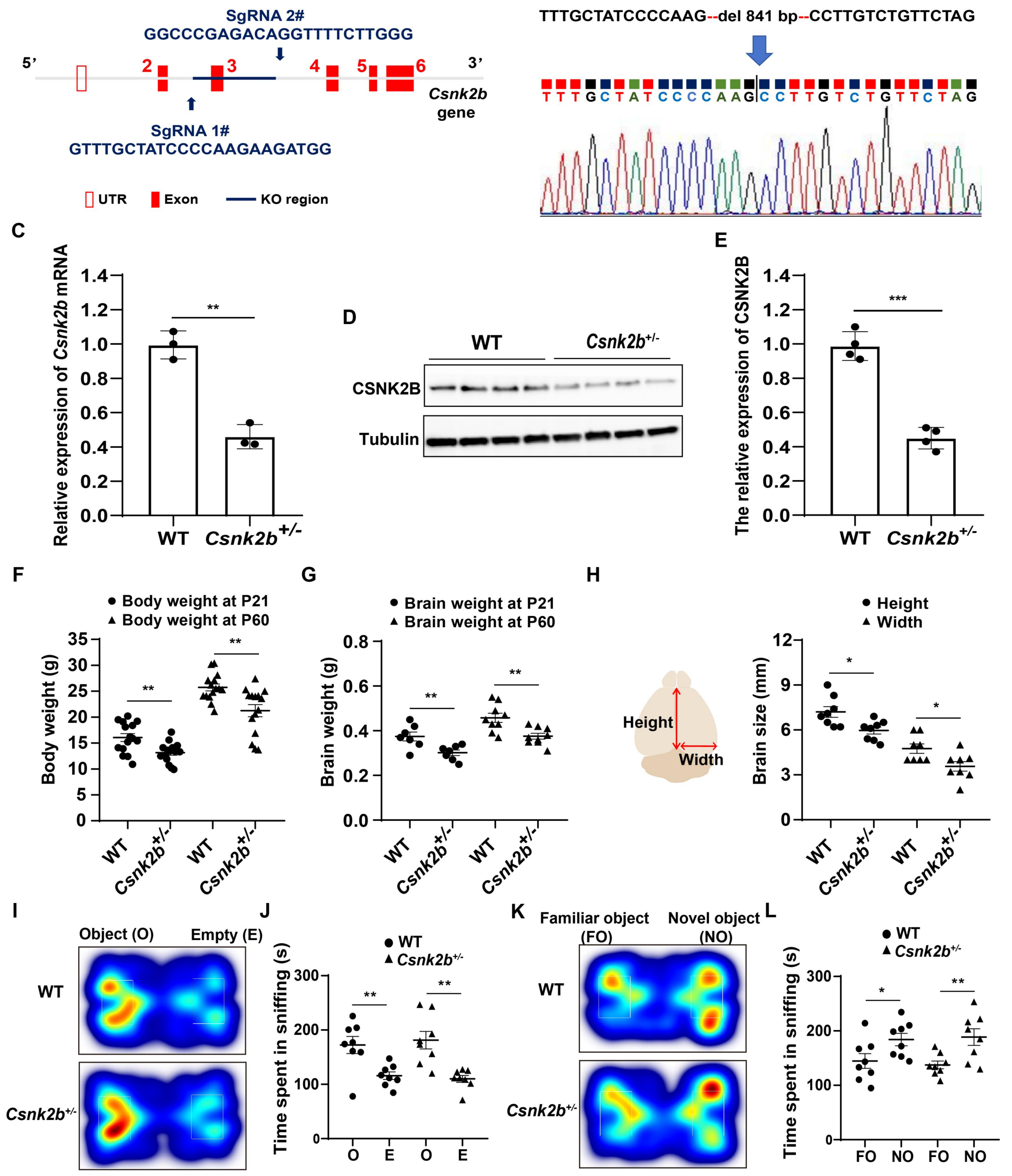


**Figure S1. Successful establishment of *Csnk2b*$^{+/-}$ mice, and their development and behavioral assessment**

(A) Specific sequences and genomic locations of the two sgRNAs utilized.

(B) Verification of an 841-nucleotide deletion in *Csnk2b*$^{+/-}$ mice through DNA sequencing.

(C) Quantitative analysis of *Csnk2b* mRNA expression in the brains of WT and

*Csnk2b*$^{+/-}$ mice (n = 3 per group).

(D) Western blot analysis of CSNK2B expression in the brains of WT and *Csnk2b*$^{+/-}$ mice (n = 4 per group).

(E) Quantification of protein expression data depicted in (D).

(F) Body weight measurements of WT and *Csnk2b*$^{+/-}$ mice at postnatal day 21 and day 60 (n = 15 per group).

(G) Brain weight measurements of WT and *Csnk2b*$^{+/-}$ mice at postnatal day 21 (n = 7 per group) and day 60 (n = 9 per group).

(H) Quantification of the height and width of the cerebral cortex in WT and *Csnk2b*$^{+/-}$ mice (n = 8 per group).

(I) Heatmap tracing of WT and *Csnk2b*$^{+/-}$ mice during the object cognition test.

(J) Time spent exploring the novel object versus empty cages (n = 8 per group).

(K) Heatmap tracing during the novel object recognition test.

(L) Quantification of time spent exploring objects (n = 8 per group).

All data are represented as mean ± SD. Statistical significance was determined using a two-tailed Student's t-test, with *$P < 0.05$, **$P < 0.01$, and ***$P < 0.001$ indicating significant differences.

Supplementary Figure 2

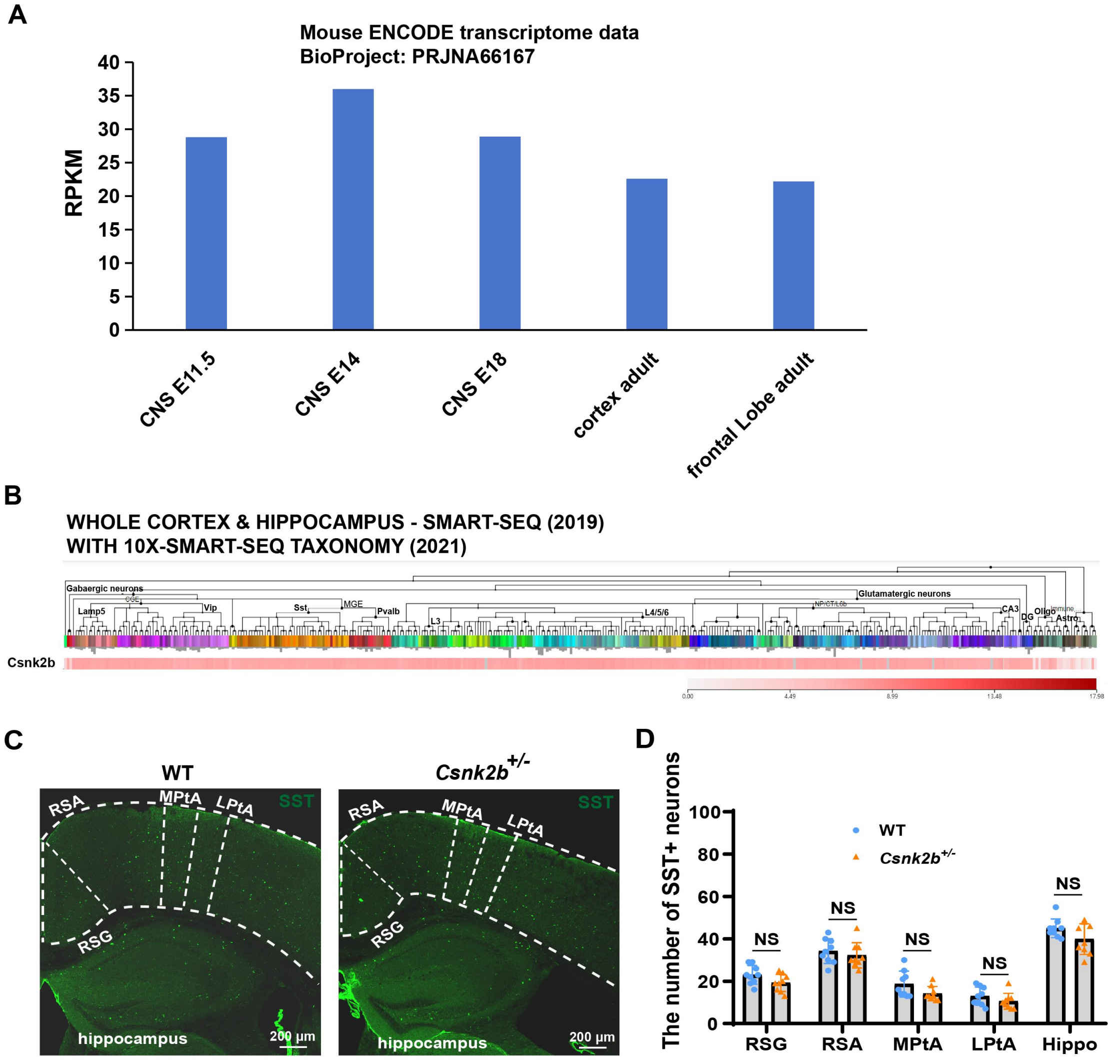


### Figure S2. Spatiotemporal expression pattern of *Csnk2b* in the mouse brain

(A) Expression levels of *Csnk2b* in the brain across different developmental stages of mice.

(B) Single-cell sequencing data of cortical and hippocampal tissues showing widespread expression of *Csnk2b* in both inhibitory and excitatory neurons.

(C) Immunostaining of SST-positive neurons in the cortex and hippocampus of WT and *Csnk2b*$^{+/-}$ mice.

(D) Quantification of SST-positive neurons in the cortex and hippocampus.

All data are represented as mean ± SD, n = 9 slices from 3 mice. Statistical significance was determined using a two-tailed Student's t-test, with $^{*}P < 0.05$, $^{**}P < 0.01$, and $^{***}P < 0.001$ indicating significant differences. NS, not significant.

Supplementary Figure 3

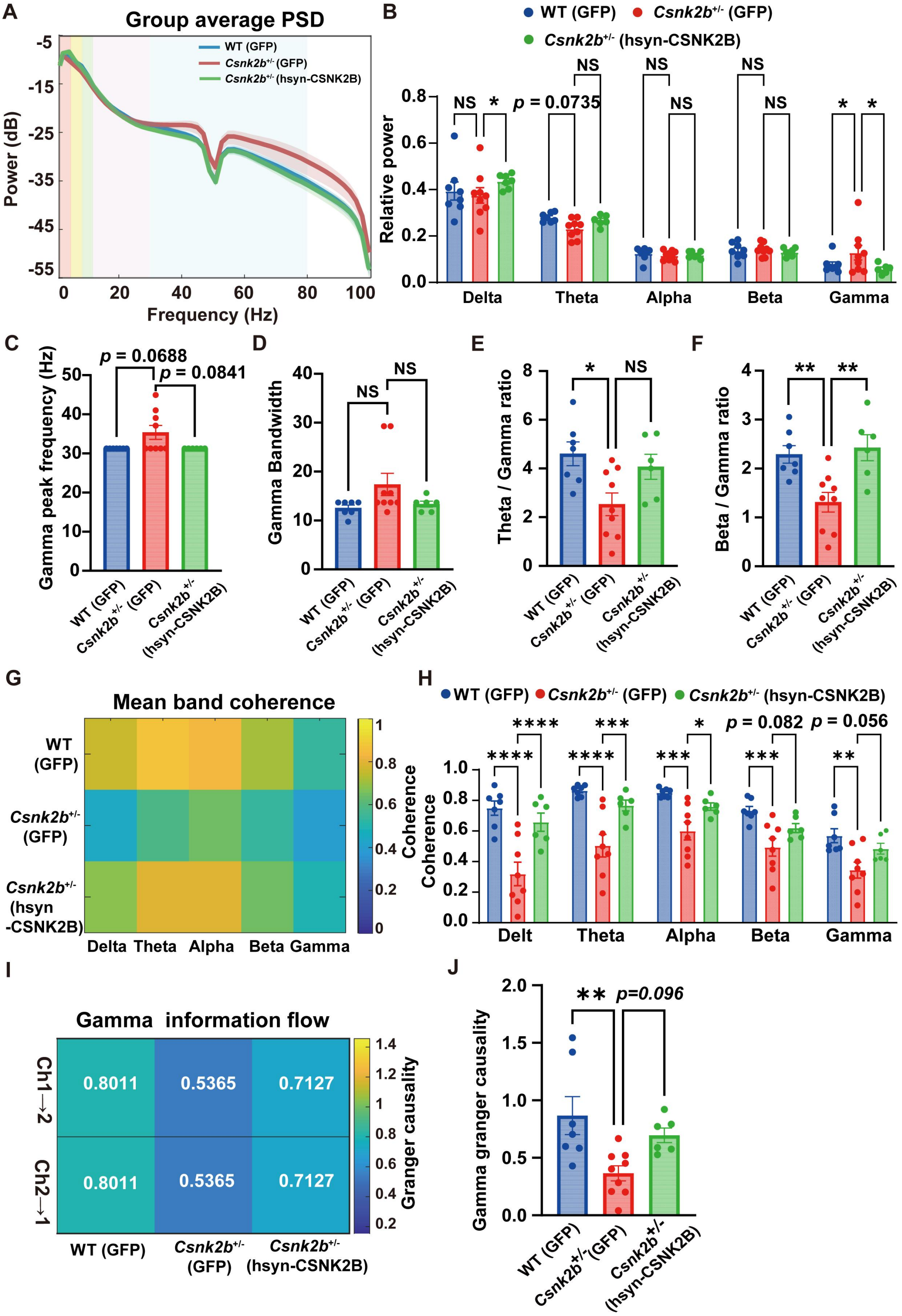


**Figure S3. AAV-PHP.eB-mediated replacement therapy stabilizes cortical E/I balance and neural network synchronization during the light (inactive) phase**

(A) Group-averaged power spectral density (PSD) of ECoG signals recorded from the mice during the light phase.

(B) Quantification of relative power across different frequency bands (Delta, Theta, Alpha, Beta, Gamma) in the three groups during the light phase.

(C) Quantification of gamma oscillation peak frequency.

(D) Quantification of gamma bandwidth.

(E) Quantification of the theta/gamma ratio in the three groups during the light phase.

(F) Quantification of the beta/gamma ratio in the three groups during the light phase.

(G) Frequency-dependent magnitude-squared coherence between bilateral PFC ECoG channels (Ch1 and Ch2) across different frequency bands in the three groups during the light phase.

(H) Quantification of mean band coherence across Delta, Theta, Alpha, Beta, and Gamma frequency bands in the three groups during the light phase.

(I) Strength of bidirectional gamma-band information flow (assessed via Granger causality) between bilateral PFC channels (Ch1→Ch2 and Ch2→Ch1) in the three groups during the light phase.

(J) Quantification of the absolute strength of bidirectional gamma-band information flow in the three groups during the light phase.

Data are expressed as mean ± SD, n = 6-12 mice for each group. Statistical analysis was performed using one way ANOVA test (C, D, E, F, J) and two-way ANOVA test (B, H). $^*P < 0.05$, $^{**}P < 0.01$, $^{***}P < 0.001$, NS (not significant).
